# Repulsion-Driven Layering in Polymer-Assisted Condensation

**DOI:** 10.64898/2026.05.08.723821

**Authors:** Arghya Majee, Holger Merlitz, Helmut Schiessel, Jens-Uwe Sommer

## Abstract

The hierarchical organization of multiphase biomolecular condensates into core-shell architectures is a fundamental problem in soft matter and biophysics. While classical explanations rely on hierarchies of interfacial tension (*γ*) between coexisting liquids, the ultralow tensions of condensates (0.1–1 *µ*N*/*m) render such hierarchies potentially fragile. We introduce a robust assembly principle based on Polymer-Assisted Condensation (PAC), in which a single polymer species dictates the entire structure. The polymer nucleates a dense core by recruiting a condensation-incompetent protein (P1). A second incompetent protein (P2), which is repelled or otherwise thermodynamically disfavored from entering the polymer-rich core, is nonetheless recruited to the interface by weak attraction to P1, forming a stable shell. This effective repulsion-driven layering operates across a wide parameter space without requiring *γ* asymmetries and yields a robust structure that is impervious to concentration fluctuations and environmental perturbations. Phase-field modeling and molecular simulations establish this mechanism and capture key features of nucleolar organization. Our work reveals a general physical pathway for encoding spatial order in soft, multicomponent fluids.

---

The spontaneous emergence of spatial order in fluid systems – from planetary atmospheres to emulsions – is a central theme in physics. In cell biology, this order is epitomized by biomolecular condensates, membrane-less organelles that compartmentalize biochemical reactions without a surrounding membrane [1–3]. These condensates embody a second level of spatial order: they not only define distinct compartments within the cell but also display internal architectures of their own [3]. Some, including the nucleolus [4], stress granules [5], paraspeckles [6, 7], and nuclear speckles [8], exhibit well-defined core–shell assemblies, whereas others, such as P granules [9] and Cajal bodies [10], display distinct though non-layered internal structuring. Understanding how such condensates generate and maintain complex spatial architectures is a central problem in soft matter and cell biology.

This problem is exemplified most prominently by the nucleolus. As the largest nuclear condensate and the site of ribosome biogenesis, it exhibits a reproducible concentric organization comprising the fibrillar center (FC), the dense fibrillar component (DFC), and the granular component (GC), with each layer enriched in distinct proteins and rRNA species [4, 11]. This persistent concentricity is particularly remarkable because condensates are generally expected to coarsen and fuse into simpler structures, not to maintain nested, multilayered architectures. Explaining the physical origin of the nucleolus’s robust organization has therefore become a major focus in the study of cellular condensates.

The prevailing physical model, drawn from emulsion thermodynamics, explains this layered organization through a hierarchy of interfacial tensions (*γ*) between distinct, coexisting liquid phases [4, 11, 12]. According to this framework, the relative surface energies of the phases with the nucleoplasm dictate their spatial arrangement, with the phase of lowest *γ* forming the outer shell. Specifically, the free-energy difference between competing morphologies scales as Δ*F* ∼ Δ*γ R*^2^, where *R* is the outer radius of the shell and Δ*γ* is the difference in interfacial tensions between configurations in which different phases (e.g., FC or DFC) form the outer interface with the surrounding phase. This model has proven powerful in rationalizing concentric organization in simple reconstituted systems.

However, biomolecular condensates operate in a regime of exceptionally low interfacial tension, typically 0.1– 1 *µ*N*/*m [13–18]. At these tensions, the capillary energy associated with a fibrillar center of radius ∼ 50 nm [19] is only of order *k*_B_*T*, indicating that such domains are only weakly stabilized by interfacial tension alone and are therefore readily perturbed by thermal fluctuations or modest variations in cellular conditions.

At the scale of the full core–shell structure (*R* ∼ 100 − 300 nm) [11, 19–21], the energetic preference between competing morphologies depends on the relevant interfacial-tension asymmetry, Δ*γ*. Even for Δ*γ* ∼ *γ*, this preference ranges from a few *k*_B_*T* for the smallest structures at the lower end of the reported tension range to a few tens of *k*_B_*T* for larger structures at the same tension. For smaller asymmetries (Δ*γ < γ*), as expected for compositionally similar core- and shell-forming phases, this scale is reduced proportionally; for example, taking Δ*γ* ∼ 0.1*γ* brings the energetic preference down to ∼ 0.3 − 3 *k*_B_*T* over *R* ∼ 100 − 300 nm at the lower end of reported tensions. As a result, multiple morphologies can be nearly degenerate in free energy, raising doubts about whether *γ*-hierarchies alone can robustly enforce concentric structure. Consistent with this view, recent theory and simulations find that when interfacial tensions are comparable, Janus-like morphologies are generic, whereas stable concentric core–shell architectures require strong asymmetry in *γ* [22].

Recent synthetic studies have shown that perturbations of rRNA processing can even invert layering under engineered conditions [23], highlighting the plasticity of nucleolar organization. Yet in vivo such inversions are not observed: nucleoli across species display robust concentricity despite environmental fluctuations and transcriptional dynamics. This points to the existence of an additional physical principle that stabilizes the canonical architecture.

Here we propose an alternative and complementary mechanism based on Polymer-Assisted Condensation (PAC) [24–26], where a central polymeric scaffold can dictate condensate architecture through a specific interaction grammar that goes beyond nucleating condensation (Fig. 1a). In this framework, a multivalent polymer (such as rRNA) acts as an architectural selector, in line with recent studies showing that RNA scaffolds can actively structure layered condensates through differential protein recruitment and segregation [7]. First, the polymer nucleates a dense core by recruiting a protein (P1) that cannot condense on its own. Second, a different protein (P2), also incompetent to phase separate alone, is energetically disfavored from the polymer-rich core yet recruited to the interface via weak attraction to P1, as encoded by the interaction scheme in Fig. 1a. This asymmetric interaction logic – polymer-mediated recruitment and exclusion – generates a stable core-shell topology through a process we term *repulsion-driven layering* (Fig. 1c).

**FIG. 1.**
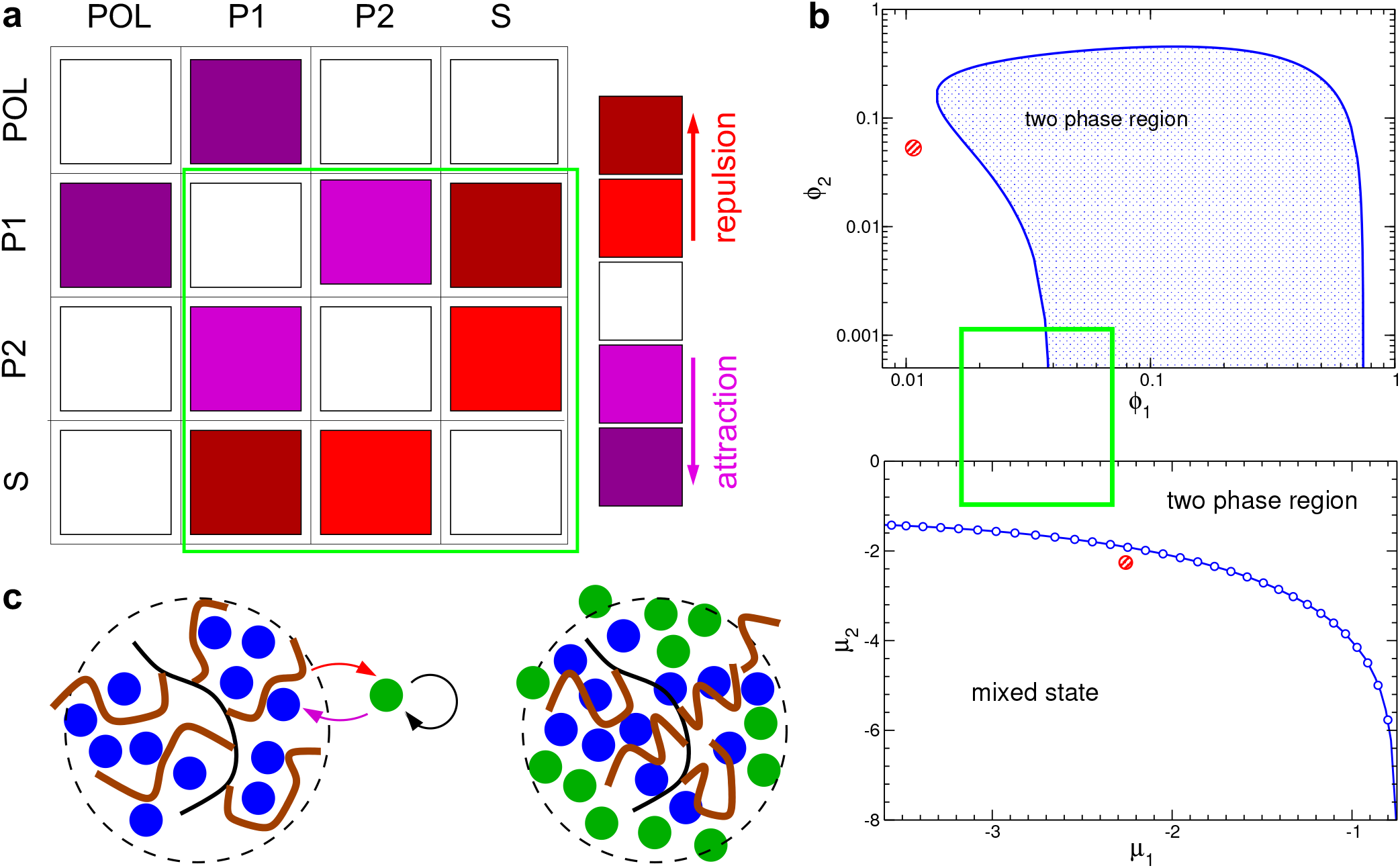
Polymer-assisted condensation (PAC) mechanism. **a** Interaction matrix between the four components: polymer (POL), protein 1 (P1), protein 2 (P2), and solvent (S). Attractive interactions are shown in purple, repulsive interactions in red, and white denotes purely excluded-volume interactions. The highlighted subset emphasizes the interaction scheme between P1, P2, and solvent that is sufficient to drive phase separation in the absence of polymer. **b** Corresponding ternary phase diagram of the subsystem (P1, P2, S) in the bulk (without polymer), shown in both concentration space (*ϕ*_1_–*ϕ*_2_, top) and chemical potential space (*µ*_1_–*µ*_2_, bottom). The shaded region indicates two-phase coexistence, while the white region corresponds to a homogeneous mixed state. The highlighted region marks the parameter regime relevant for the interaction scheme in panel (a), and the red circle indicates the state point used in the simulations. The phase diagram was obtained using the convex hull algorithm [31, 32]. For details see the method section. **c** Schematic illustration of the PAC mechanism for core–shell structure formation. The polymer (brown), representing RNA, forms an extended scaffold (e.g., a bottle-brush-like structure in the nucleolus, anchored to an underlying DNA backbone shown in black). Protein P1 (blue) is recruited to the polymer and forms a dense core via polymer-assisted condensation. Protein P2 (green), while attracted to P1, is repelled from the polymer-rich phase due to excluded-volume interactions and therefore accumulates at the interface, giving rise to a shell. The circular arrow indicates an effective self-attraction mediated by repulsion from the solvent. The results shown correspond to the parameter set *χ*_1s_ = 1.4, *χ*_2s_ = 1.0, *ε*_12_ = 0.48, and chemical potentials *µ*_1_ = *µ*_2_ = −2.258.

This pathway is fundamentally distinct from classical immiscibility, which requires the core and shell to be two pre-existing dense phases [4]. In our mechanism, the shell is not an autonomous condensate but a soluble component structured by the pre-formed core. We formalize this principle using phase-field theory and coarse-grained simulations, and show that PAC naturally explains nucleolar architecture: fibrillarin-like proteins enrich the rRNA-rich (rRNA-processing) DFC [27], whereas other nucleolar clients – such as SURF6 – can be thermodynamically disfavored from the most rRNA-enriched subphase yet retained at its interface or in the GC through multivalent recruitment by NPM1-like scaffolds [28–30]. By highlighting RNA as an active selector that both recruits and excludes, PAC provides a robust mechanism for enforcing concentric organization even in the limit of ultrasoft interfacial tensions. More generally, this framework offers a new physical principle for how polymer-mediated interactions encode spatial order in soft, multicomponent fluids.

## MEAN-FIELD CALCULATIONS

### Free energy

We model a multicomponent system comprising a long polymer, such as RNA or nucleolar organizer regions of chromatin, two protein species (P1 and P2), and an explicit solvent, all confined within a spherical domain of radius *R*. The local volume fractions of the polymer, P1, P2, and solvent are denoted by *ϕ*_p_, *ϕ*_1_, *ϕ*_2_, and *ϕ*_s_, respectively. Assuming spherical symmetry, these fields depend only on the radial coordinate *r*. The polymer is represented by an order-parameter field *ψ*(*r*) such that the local polymer volume fraction is given by *ϕ*_p_(*r*) = *ψ*^2^(*r*), and the overall (average) polymer volume fraction is denoted by Φ. We describe the polymer conformational statistics within the ground-state dominance approximation (GSDA), which is appropriate for the present setting in which the polymer is localized within the protein-rich droplet. The total free energy *F*, expressed in units of *k*_*B*_ *T*, i.e., *F* ≡ *F*_phys_*/k*_*B*_ *T*, is written as

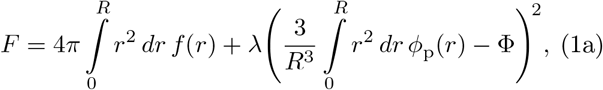

where *f* (*r*) denotes the local free-energy density. For compactness, we present it in the scaled form *νf* (*r*) (i.e., per solvent molecular volume *ν*) as

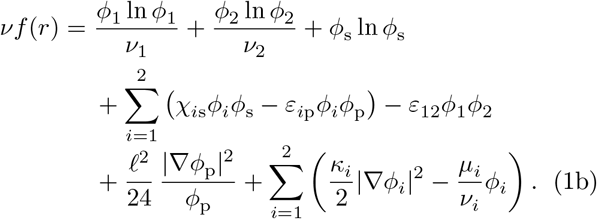

Here, *ν*_1_ and *ν*_2_ denote the dimensionless molecular-volume ratios of P1 and P2 relative to the solvent reference volume *ν*. The four volume fractions are related through the local conservation condition

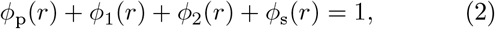

which is used to eliminate the solvent field *ϕ*_s_(*r*), thereby expressing the free energy in Eq. (1) as a functional of the three independent fields *ϕ*_p_(*r*), *ϕ*_1_(*r*), and *ϕ*_2_(*r*). The logarithmic terms in Eq. (1) represent the translational entropy of mixing of the individual components, each weighted by its molecular volume. The interaction terms proportional to *χ* and *ε* account for all pairwise enthalpic interactions between polymer, proteins, and solvent. These interactions are assumed to be short-ranged and enter the free energy as local contact terms. The elastic contribution proportional to *ℓ*^2^ arises from the chain connectivity of the polymer and corresponds to the Lifshitz entropy within the GSDA framework [33, 34]. Spatial variations in the protein concentrations are suppressed by the gradient-squared terms weighted by *κ*_*i*_, which set the energetic scale of the interfaces; higher-order and cross-gradient couplings are neglected for simplicity. The proteins are maintained at fixed chemical potentials *µ*_*i*_, coupling the system to their respective reservoirs, while the polymer is treated in the canonical ensemble with a fixed total monomer content. This constraint on the overall monomer volume fraction,

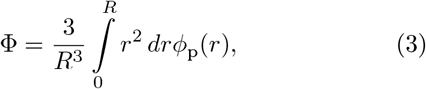

is enforced via a Lagrange multiplier *λ*, which penalizes deviations from the prescribed total monomer volume fraction. The equilibrium field profiles follow from minimizing Eq. (1) subject to the natural (Neumann) boundary conditions [35],

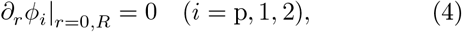

which enforce symmetry at the center and ensure that the fields reach a stationary, gradient-free state at the system boundary. Along with the global constraint in Eq. (3), these conditions determine the equilibrium spatial organization of polymer and proteins. The resulting profiles, and how they respond to changes in the model parameters, are examined below.

### Results and discussions

We begin by characterizing the baseline structure generated by the PAC mechanism and then assess how modifications of the system parameters reshape this equilibrium architecture. To connect with typical length scales in biomolecular systems, we set the unit volume to *ν* = 1 nm^3^. For comparison with molecular dynamics simulations using a Lennard–Jones model of P1, P2, and monomers in an implicit solvent, we choose *ν*_1_ = *ν*_2_ = 4.516 [36]. The polymer is assigned a Kuhn length *ℓ* = 20 nm corresponding to a semi-flexible chain. We note that these parameters can be adjusted to any particular system and the conclusions of our work do not depend on the particular parameterization.

Both proteins tend to demix from the solvent, with Flory parameters *χ*_1s_ = 1.4 and *χ*_2s_ = 1.0. However, these interactions alone are insufficient to drive bulk phase separation under our conditions. For a given *χ*, coexistence occurs only at a specific chemical potential *µ* = *µ*_coex_(*χ*). We deliberately choose *µ*_1_ = *µ*_2_ = −2.258, which lies below this coexistence value, such that the homogeneous dilute phase remains the only stable bulk state (Fig. 1b). Under these conditions, the system remains in the single-phase regime of the ternary mixture, such that neither individual nor cooperative phase separation of the proteins occurs in the absence of the polymer. Condensation can nevertheless emerge through the PAC mechanism (Fig. 1c), in which the polymer acts as an extended, attractive scaffold that stabilizes local enrichment of protein [24–26]. We choose interaction strengths *ε*_1p_ = 6.0, *ε*_2p_ = 0 to ensure that only protein 1 (P1) can form a polymer-mediated condensate via PAC, while protein 2 (P2) remains below the threshold required for phase separation. In addition, the two protein components also experience a weak mutual attraction, quantified by *ε*_12_ = 0.48. The interfacial properties of both proteins are governed by equal gradient coefficients *κ*_1_ = *κ*_2_ = 10 nm^2^, which produce diffuse interfaces several nanometers wide. These values are consistent with experimental observations of nanometer-scale surface layers in biomolecular condensates [37], as well as with diffuse coacervate interfaces reported in simulations [38] and fall within the typical parameter ranges used in Cahn–Hilliard–type models of phase-separated biomolecular systems [39–41].

Figure 2 presents the steady-state concentration fields of this ternary polymer–protein system. Panel a shows a two-dimensional polar map of the polymer volume fraction *ϕ*_p_, displayed using a sequential yellow–brown color scale. Overlaid in green is the distribution of P2, *ϕ*_2_, rendered with a graded transparency to preserve the visibility of the underlying polymer scaffold. Panel b displays the corresponding spatial distribution of P1, *ϕ*_1_, shown separately in blue for clarity. Panel c shows the one-dimensional radial profiles *ϕ*_p_, *ϕ*_1_, and *ϕ*_2_, enabling direct quantitative comparison across components.

**FIG. 2.**
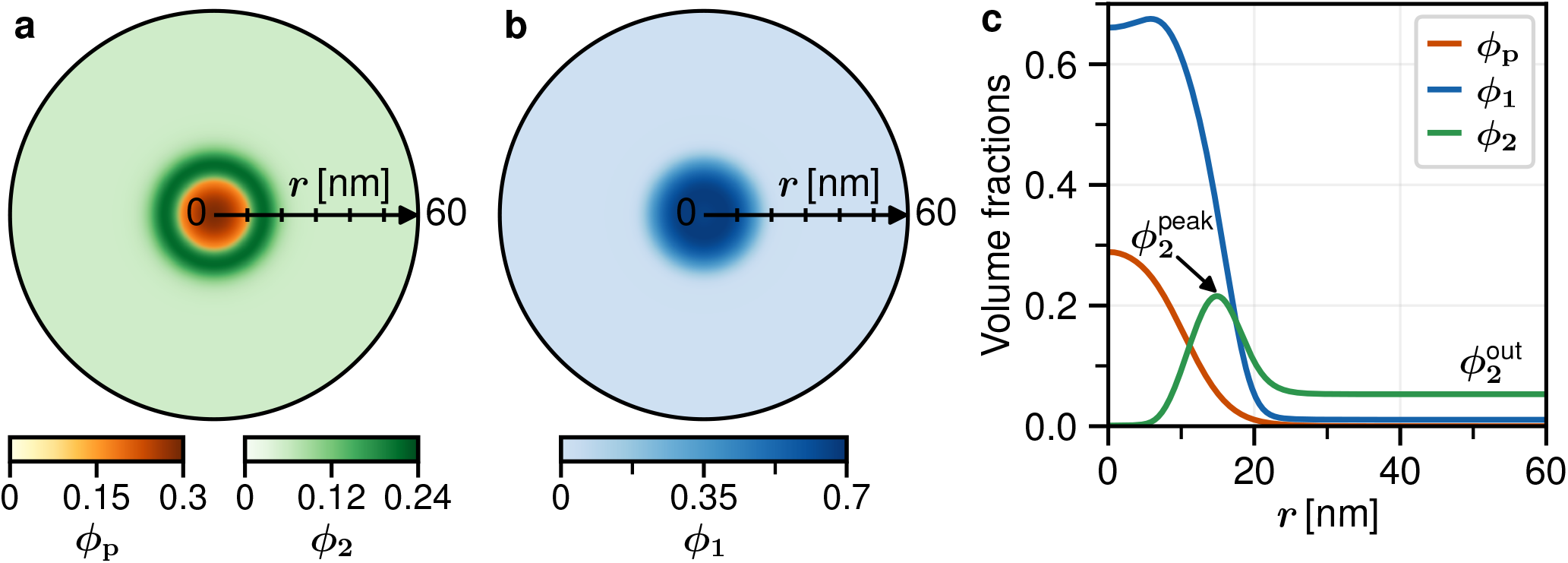
Stable core–shell architecture arising from polymer-assisted condensation (PAC) and repulsion-driven layering. **a** Polar map of the polymer volume fraction *ϕ*_p_(*r*) = *ψ*^2^(*r*), encoded by a sequential yellow–brown color scale and overlaid with the P2 concentration *ϕ*_2_ (green). The inner core corresponds to the RNA-like polymer scaffold, where polymer–P1 attraction enriches the polymer region and suppresses P2. The surrounding shell is formed by P2, whose spatial distribution reflects the balance between polymer-mediated exclusion and weak attraction to P1 at the core–shell interface. **b** Polar map of the P1 concentration *ϕ*_1_ (blue), shown on the same radial grid and colour-scaled independently from panel (**a**). The polymer-enriched interior coincides with a high concentration of P1, consistent with polymer-mediated recruitment. Horizontal colour bars beneath panels (**a**) and (**b**) indicate the colormap ranges for *ϕ*_p_(*r*), *ϕ*_1_(*r*), and *ϕ*_2_(*r*). **c** Radial profiles of *ϕ*_p_(*r*), *ϕ*_1_(*r*), and *ϕ*_2_(*r*) that underlie the two-dimensional maps. The profiles illustrate the PAC mechanism: polymer recruitment of P1 produces a dense core, while polymer exclusion combined with P1–P2 attraction localizes P2 to the outer shell, establishing a robust core–shell structure. The results shown correspond to the parameter set *χ*_1s_ = 1.4, *χ*_2s_ = 1.0, *ε*_1p_ = 6.0, *ε*_12_ = 0.48, chemical potentials *µ*_1_ = *µ*_2_ = − 2.258, surface parameters *κ*_1_ = *κ*_2_ = 10 nm^2^. Importantly, the chosen chemical potentials place both protein species outside their bulk coexistence regime, such that neither P1 nor P2 undergoes phase separation in the absence of the polymer; the core–shell architecture therefore arises entirely from polymer-mediated recruitment and exclusion (Fig. 1). The maximum shell concentration of protein 2, 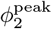, and its dilute-phase value far from the condensate, 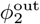, are indicated and used to quantify shell enrichment in subsequent analysis.

These profiles reveal a condensate with a well-defined core–shell architecture. A dense polymer–P1 condensate forms at the center, *r* = 0, driven by the polymer–protein interaction. Protein P2, which lacks affinity for the polymer (*ε*_2p_ = 0), is excluded from the condensed core. Nevertheless, its weak attraction to protein 1 prevents complete dispersion into the dilute phase, leading instead to its accumulation at the condensate periphery. There, it forms a P2-rich corona with a maximum volume fraction 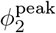, which is approximately four times higher than its dilute-phase bulk concentration 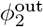. This spatial organization arises from the interplay between selective polymer binding and differential interspecies interactions, illustrating how modest variations in molecular affinity can give rise to complex internal architectures within biomolecular condensates.

Next, we examine how the properties of the P2-rich shell depend on system parameters. We characterize the shell using two quantities: the enrichment of protein 2, quantified by the ratio 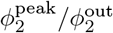 (see Fig. 2c for a visual illustration), and its geometry, described by the inner and outer radii *R*_in_ and *R*_out_, as well as the shell thickness *δR* = *R*_out_ − *R*_in_. These radii are extracted from the radial profile of *ϕ*_2_(*r*) using a standard half-maximum criterion (see Methods).

Figure 3 summarizes how these quantities vary with three key parameters: the heterotypic interaction strength *ε*_12_ between proteins 1 and 2, the chemical potential *µ*_2_ of protein 2, and the gradient coefficient *κ*_2_. Increasing *ε*_12_ enhances the preferential accumulation of protein 2 at the condensate periphery, leading to an increase in the enrichment ratio 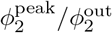 (Fig. 3a). This behavior arises because increasing *ε*_12_ strengthens the attraction between P1 and P2, lowering the free energy of protein 2 in the interfacial region and promoting its accumulation at the condensate boundary.

**FIG. 3.**
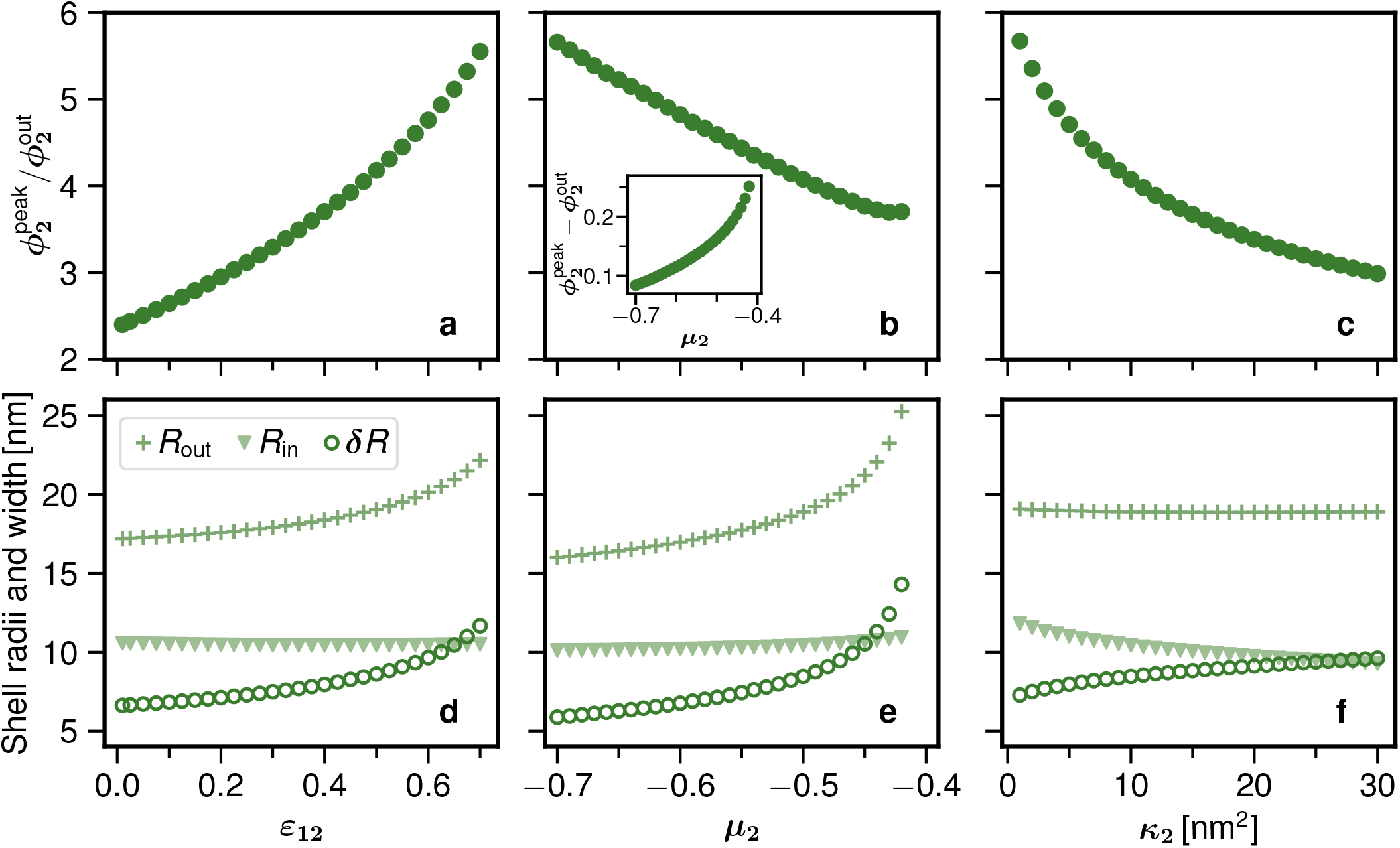
Mean-field prediction of shell composition and geometry in Polymer-Assisted Condensation. **a–c** Peak shell enrichment of protein 2, quantified as 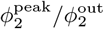, as a function of (**a**) the P1–P2 interaction strength *ε*_12_, (**b**) the chemical potential of protein 2, *µ*_2_, and (**c**) the gradient coefficient *κ*_2_. Increasing *ε*_12_ enhances the preferential accumulation of protein 2 at the condensate periphery, leading to a higher enrichment ratio. In contrast, increasing *µ*_2_ reduces the enrichment ratio, reflecting a concurrent increase of protein 2 in both the shell and the surrounding dilute phase. Increasing *κ*_2_ suppresses sharp spatial variations in *ϕ*_2_ and reduces shell enrichment. Inset in (**b**) shows the corresponding absolute enrichment, 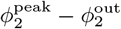, which increases with *µ*_2_, indicating that the total accumulation of protein 2 at the interface grows even as the relative contrast decreases. **d–f** Corresponding variations of the inner shell radius *R*_in_, outer shell radius *R*_out_, and shell thickness *δR* = *R*_out_ − *R*_in_ as defined in Eqs. (6) and (7). Increasing *ε*_12_ (**d**) or *µ*_2_ (**e**) thickens the shell primarily through an outward shift of *R*_out_, while *R*_in_ remains approximately fixed, reflecting that the polymer–P1–rich core remains thermodynamically unfavorable for protein 2. In contrast, increasing *κ*_2_ (**f** ) broadens the shell predominantly by shifting *R*_in_ inward at nearly constant *R*_out_, consistent with the reduced energetic cost of expanding the diffuse layer at smaller radii. Together, these results establish a continuum-level description of repulsion-driven layering within the PAC framework. Unless otherwise stated, the parameters *µ*_1_ = *µ*_2_ = − 2.258 and *ε*_12_ = 0.48 are used.

In contrast, increasing *µ*_2_ reduces the enrichment ratio (Fig. 3b), even though the absolute accumulation of protein 2 at the interface increases. This behavior arises because a higher chemical potential elevates the concentration of protein 2 throughout the system, leading to a concurrent increase in both 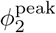 and 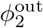, with the latter increasing more strongly. As a result, the relative contrast between the shell and the surrounding phase decreases. Increasing *κ*_2_ suppresses sharp spatial variations in *ϕ*_2_, leading to a reduced enrichment ratio (Fig. 3c).

The corresponding changes in shell geometry are shown in Fig. 3d–f. Increasing *ε*_12_ or *µ*_2_ thickens the shell primarily through an outward shift of the outer radius *R*_out_, while the inner radius *R*_in_ remains approximately unchanged. This behavior reflects that the inner boundary is set by the polymer–P1–rich core, which remains thermodynamically unfavorable for P2, whereas the interfacial region provides the lowest free-energy environment for protein 2. By contrast, increasing the gradient coefficient *κ*_2_ broadens the shell predominantly by shifting the inner radius inward at nearly constant outer radius. This asymmetry arises because spreading the P2 profile toward smaller radii reduces the area of the inner interface, whereas expanding the outer radius would increase the interfacial area and therefore incur a larger gradient-energy penalty. Together, these results establish a continuum-level picture of repulsion-driven layering within the PAC framework, motivating a microscopic validation using particle-based simulations.

## MOLECULAR DYNAMICS SIMULATIONS

### Simulation model and setup

To complement our field-theoretic framework and to directly interrogate the microscopic underpinnings of PAC, we constructed a coarse-grained molecular dynamics model that captures the essential interactions among a flexible polymer and two protein-like components. All simulations were performed in reduced Lennard–Jones (LJ) units, with the LJ energy scale *ε*, length scale *σ*, and Boltzmann constant *k*_*B*_ set to unity (*i*.*e*., *ε* = *σ* = *k*_*B*_ = 1). The polymer is represented as a FENE bead–spring chain of length *N* = 800 monomers in a cubic simulation box with periodic boundary conditions in all directions. Two types of monomeric cosolvents, corresponding to the condensation-incompetent proteins P1 and P2, populate the surrounding environment.

Nonbonded interactions between all species are modeled using truncated and shifted Lennard–Jones potentials. The interaction strengths are specified in units of *ε* and are denoted by 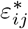, allowing independent control over polymer–cosolvent and cosolvent–cosolvent affinities. Unless otherwise stated, the interaction matrix is chosen such that polymer–polymer and polymer–P2 interactions are purely repulsive, polymer–P1 interactions are attractive, and P1–P1, P2–P2, and P1–P2 interactions are weakly attractive. The chemical potentials of both cosolvent species are fixed. (See Methods for numerical values.) This choice encodes the three core ingredients of PAC, namely attractive polymer–P1 interactions, repulsive polymer–P2 interactions, and weak attraction between P1 and P2.

To allow the local concentrations of P1 and P2 to self-adjust in response to these interactions, both cosolvent species are coupled to particle reservoirs via grand-canonical Monte Carlo insertion–deletion moves at fixed reduced temperature *T*^∗^ = *k*_*B*_ *T/ε* = 1. The chemical potentials of the cosolvents are specified in reduced form as 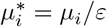, enabling dynamic recruitment or expulsion of P1 and P2 without imposing fixed stoichiometries.

### Simulation results

To enable a qualitative comparison with the mean-field predictions, we systematically vary three control parameters in the molecular dynamics simulations: the P1–P2 interaction strength 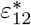, the chemical potential of protein 2, 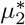, and the polymer–P2 interaction strength 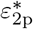, all expressed in units of the Lennard–Jones energy scale *ε*. As a reference case, we consider purely repulsive P1–P2 interactions, implemented by truncating the Lennard–Jones potential at its minimum such that no attractive tail is present. The resulting steady-state concentration profiles are shown in Fig. 4.

**FIG. 4.**
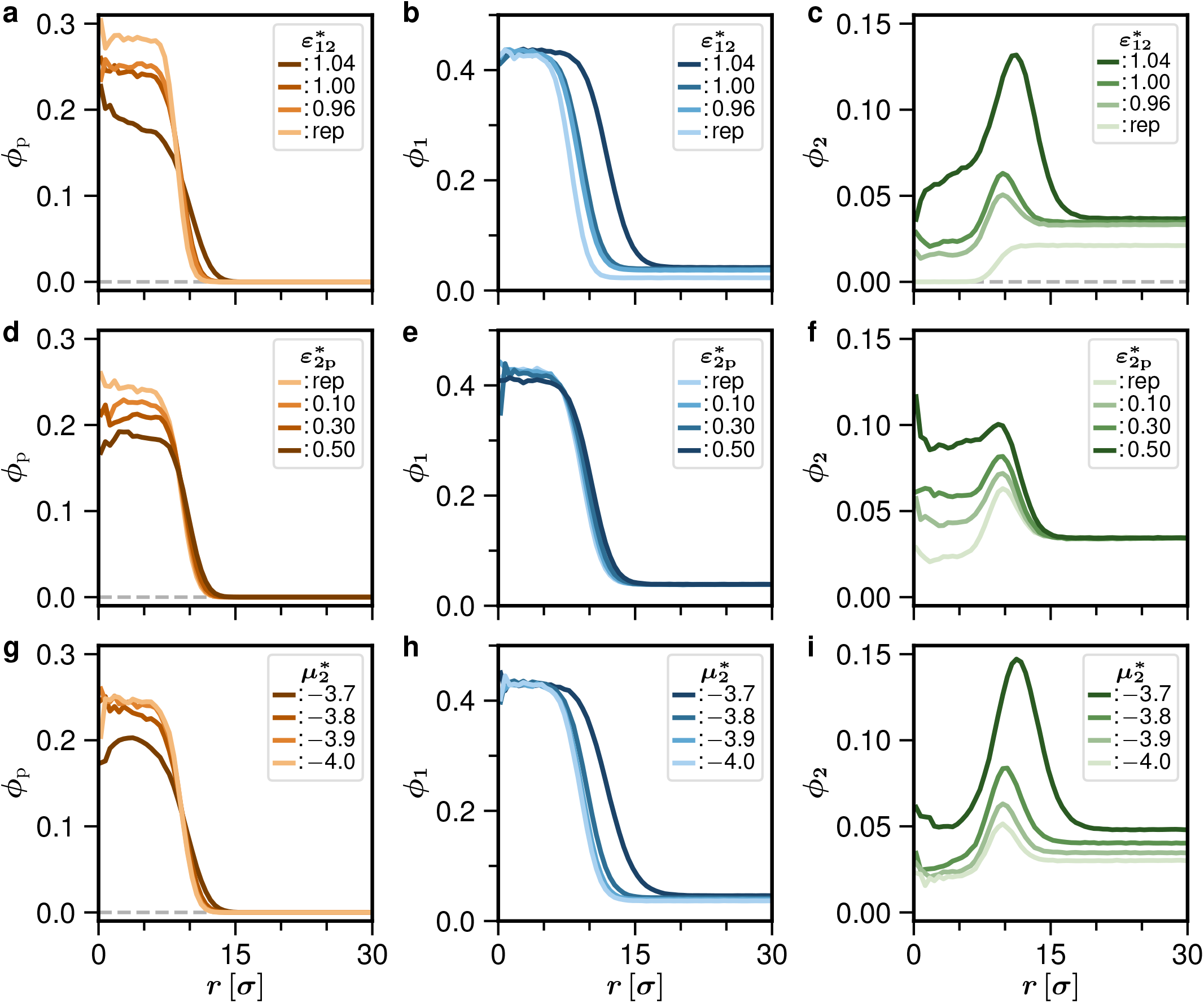
Molecular dynamics simulations confirm repulsion-driven layering at the particle level. **a–c** Radially averaged volume-fraction profiles of polymer, protein 1 (P1), and protein 2 (P2) as the P1–P2 interaction strength 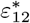 is increased from a purely repulsive reference case to weakly attractive interactions. Strengthening 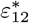 leads to enhanced recruitment of P2 to the condensate periphery, resulting in both a higher peak concentration within the shell and an increased shell thickness, while the polymer–P1–rich core remains largely unchanged. **d–f** Corresponding profiles as the polymer–P2 interaction strength 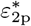 is varied from a purely repulsive reference case to attractive interactions. Increasing 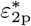 promotes incorporation of P2 into the condensate interior, while the dilute-phase concentration remains essentially unchanged. The increased P2 population is accommodated by a slight reduction of polymer density within the condensate and a corresponding modest expansion of the droplet, consistent with conservation of polymer content. **g–i** Profiles obtained upon increasing the chemical potential 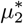 of protein 2. Raising 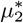 increases the reservoir concentration of P2 and leads to a pronounced accumulation at the interface, again producing a thicker shell without perturbing the core structure. In both cases, the increased P2 population is accommodated by a modest redistribution of local volume fractions rather than a restructuring of the polymer–P1 core. Unless otherwise stated, parameters are fixed to 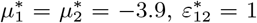, and a repulsive polymer–P2 interaction 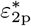. These particle-based results qualitatively reproduce the mean-field predictions and demonstrate that repulsion-driven layering emerges robustly at the microscopic level.

Figures 4a–c illustrate the system response as the P1–P2 interaction is tuned from the purely repulsive reference case to weakly attractive interactions with increasing 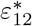. In agreement with the field-theoretic results, strengthening the P1–P2 attraction leads to enhanced recruitment of protein 2 to the condensate periphery, manifested as both an increased peak concentration within the shell and an increased shell thickness. Notably, the size of the polymer-rich core remains largely unchanged across this range of 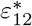. As a consequence, shell growth occurs predominantly through an outward expansion of the condensate, as confirmed by the P1 volume fraction profiles in Fig. 4b. Accommodating the increased P2 population is accompanied by a modest reduction in the local polymer volume fraction, reflecting volume fraction redistribution rather than a restructuring of the core.

The effect of polymer–P2 interactions is shown in Fig. 4d–f, where the polymer–P2 interaction strength 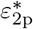 is varied from a purely repulsive reference case to attractive interactions. Increasing 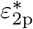 promotes the incorporation of protein 2 into the condensate interior, leading to a higher P2 concentration within the droplet while leaving the dilute-phase concentration largely unchanged. This redistribution is accompanied by a slight decrease in the polymer volume fraction within the condensate and a corresponding modest expansion of the droplet, consistent with conservation of polymer content. Despite this redistribution, the overall structure of the polymer–P1–rich core remains intact.

A similar but distinct phenomenology is observed upon varying the chemical potential 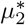, as shown in Fig. 4g–i. Increasing 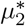 raises the reservoir concentration of protein 2 and leads to a corresponding increase of its volume fraction throughout the system, with the most pronounced accumulation occurring within the interfacial shell. This results in both a higher peak concentration and a thicker shell, while the P1–rich condensate core remains essentially unaffected. As in the case of varying 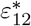, the increased P2 population is accommodated by a modest redistribution of local volume fractions, without altering the overall organization of the polymer–P1 core.

To quantify these trends, we extract the shell enrichment and geometry from the simulation data (Fig. 5). Increasing 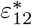 produces a pronounced increase in the enrichment ratio 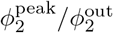, Fig. 5a, reflecting stronger P1–P2 attraction that stabilizes protein 2 at the condensate interface. Increasing 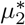 leads to a weaker increase, Fig. 5b, arising from a global rise in protein 2 concentration combined with preferential accumulation at the interface. A similar increase is observed upon increasing 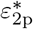, Fig. 5a, which enhances the partitioning of protein 2 into the condensate without significantly altering its dilute-phase concentration.

**FIG. 5.**
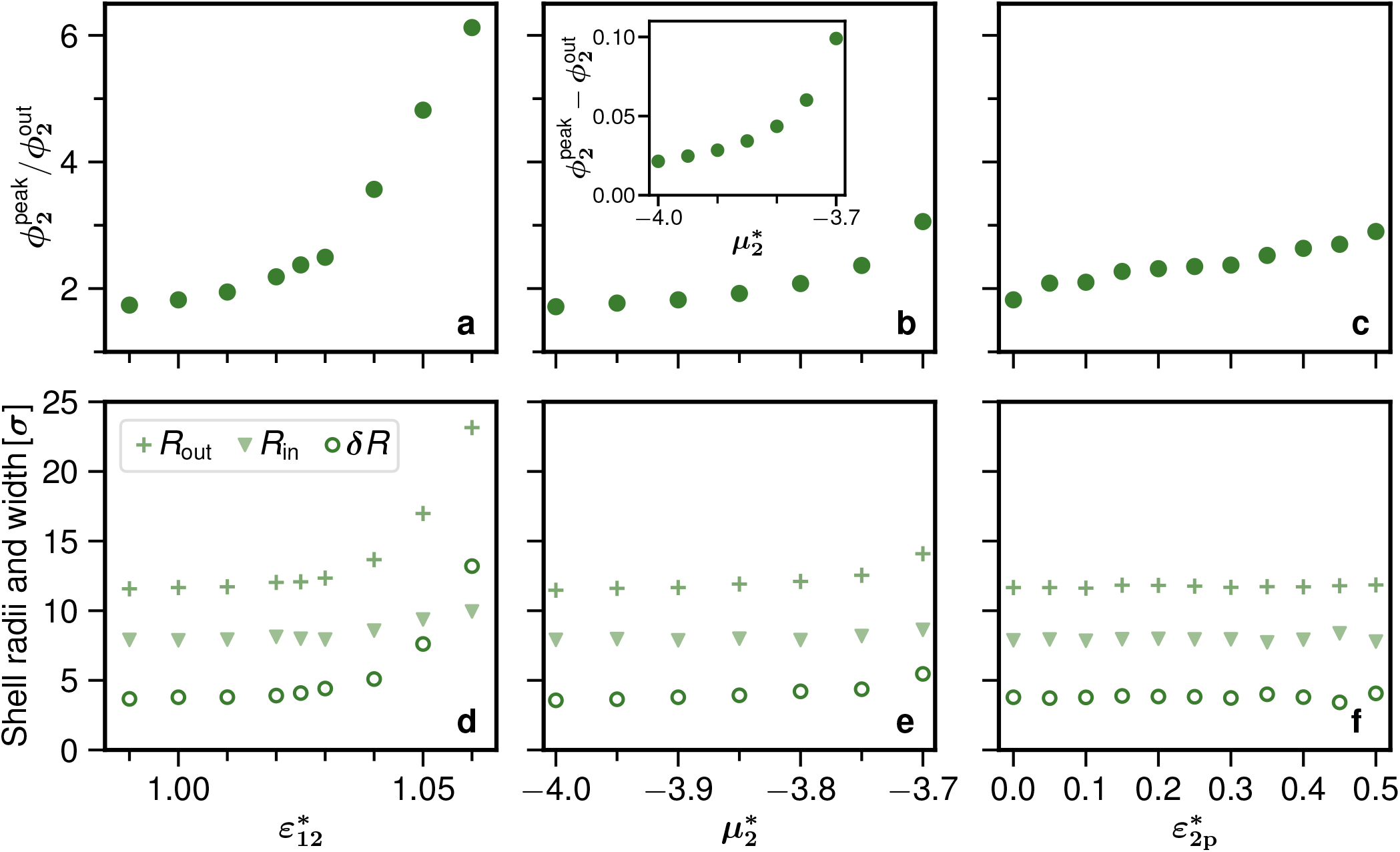
Quantitative characterization of shell enrichment and geometry in molecular dynamics simulations. **a–c** Peak shell enrichment of protein 2, quantified as the ratio 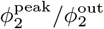, as a function of (**a**) the P1–P2 interaction strength 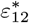, (**b**) the chemical potential 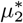, and (**c**) the polymer–P2 interaction strength 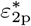. Increasing each of these parameters enhances the accumulation of protein 2 at the condensate periphery, leading to a higher enrichment ratio. In the case of increasing 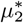, this reflects a combined increase of protein 2 in both the shell and the surrounding phase, with a stronger relative increase at the interface. **d–f** Corresponding variations of the inner shell radius *R*_in_, outer shell radius *R*_out_, and shell thickness *δR* = *R*^out^ − *R*^in^ (defined in Eqs. (6) and (7)) as functions of (**d**) 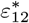, (**e**) 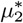, and (**f** ) 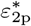. Increasing 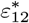 leads to a pronounced growth of the shell, primarily through an outward shift of *R*_out_, while *R*_in_ changes more weakly. A similar but less pronounced trend is observed upon increasing 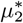. In contrast, increasing 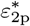 leaves both shell radii and thickness nearly unchanged, consistent with polymer–P2 attraction promoting partitioning of protein 2 into the condensate interior without altering the overall shell geometry. As in Fig. 4, unless otherwise stated, parameters are fixed to 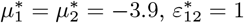, and a repulsive polymer–P2 interaction 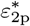. Together, these results demonstrate that distinct interaction parameters control shell enrichment and geometry in different ways, in agreement with the mean-field predictions.

The corresponding changes in shell geometry are shown in Fig. 5d–f. Increasing 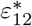 leads to a significant growth of the shell, primarily through an outward shift of *R*_out_, while *R*_in_ changes more weakly. A similar but less pronounced trend is observed upon increasing 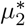. In contrast, increasing 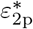 leaves both shell radii and thickness nearly unchanged, indicating that polymer–P2 attraction predominantly redistributes protein 2 within the condensate without altering the overall shell geometry.

Taken together, the field-theoretic and molecular simulations demonstrate that repulsion-driven layering is a robust and generic consequence of Polymer-Assisted Condensation, persisting across modeling scales.

## CONCLUSIONS

In this work, we have identified a robust physical mechanism by which complex internal architectures can arise in multicomponent biomolecular condensates without relying on hierarchies of interfacial tension. Within the PAC framework, a single polymeric scaffold plays a dual architectural role: it selectively recruits certain proteins to nucleate a dense core while simultaneously disfavoring others from the interior. When combined with weak heterotypic attractions, this asymmetry naturally drives interfacial localization and stabilizes a core–shell organization through a process we term repulsion-driven layering. Importantly, in this mechanism the shell does not constitute an autonomous condensed phase, but instead emerges as a soluble component structured around a polymer-driven core. Let us reemphasize the guiding principle in PAC: All protein species display a miscibility gap but their concentration is outside of it. This prevents spontaneous droplet formation everywhere in the cyto- or nucleoplasm. In the presence of a polymer that selectively attracts at least one protein species, local condensation is induced within the spatial extent of the polymer conformations. Thus condensates are controlled in space. The locally condensed phase can further interact with other components or environmental factors such as membranes and surfaces. In particular co-condensation with a second species is frustrated by the presence of the polymer which leads to spatial organization such as core-shell structures reported in this work. Notably, this mechanism does not depend on surface tension effects but is stabilized by the conformational flexibility (entropy) of the polymer. By retreating from parts of the condensate in favor of co-condensation between the two protein species, the polymer is governing and stabilizing the morphology of the co-condensate. As a direct consequence, the core–shell condensate is not self-sustaining: removal of the polymer scaffold leads to its complete dissolution.

Using a phase-field description, we showed how shell enrichment and thickness respond to variations in interaction strengths, chemical potentials, and gradient penalties. This also demonstrate the robustness of the core-shell morphology with respect to parameter variation. Coarse-grained molecular dynamics simulations confirm that the same organizational logic persists at the particle level, even when correlations and thermal fluctuations are naturally incorporated at the microscopic level. The qualitative agreement across continuum and particle-based descriptions underscores the generality of the mechanism and demonstrates that repulsion-driven layering is the rule across modeling frameworks, rather than an exception or an artifact of a particular level of description.

More broadly, current biophysical pictures of nucleolar organization are best viewed as complementary rather than mutually exclusive. In the chromatin–crosslinking paradigm [42], rDNA-bearing chromatin acts as an organizing scaffold whose transient bridging interactions can drive spatial segregation and regulate viscoelastic properties. In the condensate paradigm, as developed here, RNA–protein mixtures form multiphase-separated materials whose internal architecture emerges from interaction asymmetries, with RNA acting as both recruiter and excluder of client species. Within this framework, Polymer-Assisted Condensation provides a mechanism for stabilizing core–shell order even in the limit of ultrasoft interfacial tensions.

A natural unifying picture is that nucleolar organization arises from a viscoelastic, multicomponent condensate embedded within and mechanically coupled to a dynamically cross-linked chromatin network. In such a scenario, crosslink kinetics may impose spatial and temporal constraints on local phase behavior, while condensate formation and transcriptional activity feed back on chromatin organization.

The PAC framework further highlights the key role of large biomacromolecules such as RNA or chromatin sequences as architectural selectors in biomolecular condensates. Rather than serving as passive scaffolds, these polymers can simultaneously nucleate condensation-incompetent proteins and exclude others, while undergoing conformational changes such as compaction, thereby encoding spatial organization through asymmetric interaction logic. This perspective is particularly consistent with the biochemical composition of the nucleolus, which is enriched in weakly interacting client proteins whose localization depends on rRNA-mediated recruitment. We anticipate that similar polymer-mediated principles may operate in other RNA/DNA-rich condensates, providing a general route by which spatial order can be established and maintained in soft, multicomponent fluids despite ultrasoft interfaces and biological fluctuations, with potential relevance extending beyond nuclear organization to synthetic condensates and engineered soft materials.

## METHODS

### Equilibrium phase diagram of the protein subsystem

The bulk solution outside of the polymer can be described by the homogeneous free energy density according to Eq. (1b) in units of *k*_B_*T* and the molecular volume of the solvent:

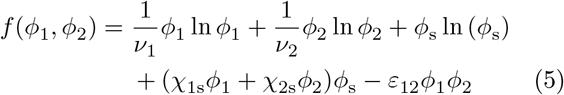

with *ϕ*_1_ + *ϕ*_2_ + *ϕ*_s_ = 1. The lower convex hull is then obtained from the discretized free-energy density over the ternary phase space. The calculation of the binodals is based on an analysis of the symmetry of the triangles resulting from the convex-hull algorithm [31]. The upper part of Fig. 1b corresponds to the parameter choice used in the main text for the bulk: *ν*_1_ = *ν*_2_ = 4.516 (which provides the best approximation to the Lennard-Jones model with implicit solvent), *χ*_1s_ = 1.5, *χ*_2s_ = 1.0, and *ε*_12_ = 0.48. Note that in the limit *ϕ*_2_ → 0, the binding protein (*ϕ*_1_) exhibits a miscibility gap, while the non-binding protein lies outside the two-phase region in the absence of the binding protein (*χ*_2_ *< χ*_*c*_ ≃ 1.0813). The attraction between the two proteins leads to co-condensation with increasing concentration of the non-binding species in the region 0.013 *< ϕ*_1_ *<* 0.038. However, in this work, we consider a bulk state outside the coexistence region. In the space of chemical potentials, the coexistence region collapses to a single line, as shown in the lower part of Fig. 1b.

The numerical calculations were performed using, in particular, the *ConvexHull* class from the Python *scipy* package. Parts of the code (optimization routines/refactoring) were developed with the assistance of an AI language model (OpenAI ChatGPT, GPT-5.1) and were subsequently reviewed, tested, and adapted by the authors (JUS).

### Mean-field calculations

The free energy in Eq. (1) is minimized numerically on a 1D discretized domain using a gradient descent algorithm. The integral has been evaluated using a trapezoidal rule and the derivatives have been represented within the finite difference approximation. The implementation is carried out using a custom C++ code.

The minimization is performed with respect to the spatially varying volume fractions *ϕ*_p_, *ϕ*_1_, and *ϕ*_2_. For numerical convenience, the polymer volume fraction is represented by an auxiliary variable *ψ* defined through *ϕ*_p_ = *ψ*^2^, such that the free energy is minimized in terms of *ψ, ϕ*_1_, and *ϕ*_2_.

To improve numerical stability and ensure that all volume fractions remain within their physical bounds, the minimization is carried out in terms of transformed variables *x, y*, and *z*. Specifically, the polymer field is parametrized as *ψ* = tanh(*x*), such that *ϕ*_p_ = *ψ*^2^ ∈ [0, 1], while the remaining volume fraction 1 − *ϕ*_p_ is distributed between the two protein species according to

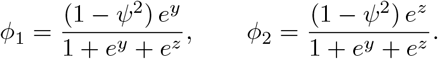

This parametrization ensures positivity of all components and automatically enforces the local volume-fraction constraint, Eq. (2). Separate step sizes, *τ*_*x*_, *τ*_*y*_ and *τ*_*z*_, are employed for the three fields to account for their different relaxation scales. The iterative updates are given by

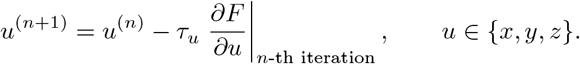

In practice, the derivatives are evaluated using all fields at their values at iteration *n*, i.e., using *x*^(*n*)^, *y*^(*n*)^, *z*^(*n*)^ . Convergence is assessed using two complementary criteria. First, the relative change in free energy between successive iterations is monitored and required to fall below a prescribed tolerance, typically approaching the limits of floating-point precision, such that no further decrease in free energy is observed. Second, the maximal update magnitude of the fields between iterations is monitored and required to become negligible. In practice, the minimization is terminated only when both conditions are satisfied. To avoid convergence to metastable states, hundreds of randomized initial profiles are tested, and the configuration yielding the lowest free energy is selected. To enforce the monomer volume fraction conservation constraint in Eq. (3), we employ a high *λ*-value (see Eq. (1)), ensuring that any deviation from the constraint incurs a significant energy penalty, thereby guaranteeing its satisfaction.

From the converged radial profiles, we extract the geometric characteristics of the P2-rich shell. The shell enrichment is quantified by the peak value 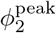 relative to the dilute-phase concentration 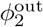. The shell boundaries are defined by the inner and outer radii, *R*_in_ and *R*_out_, at which the P2 volume fraction reaches half of its excess above the corresponding inner and outer reference concentrations. Specifically, these radii satisfy

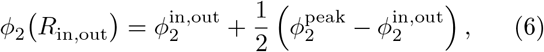

where 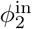 and 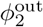 denote the P2 concentrations on the inner and outer sides of the shell, respectively. The shell thickness is then defined as

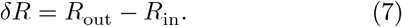

### Molecular dynamics simulations

To complement the field-theoretic description, we performed coarse-grained molecular dynamics simulations using the LAMMPS package [43–45]. All simulations were carried out in reduced Lennard–Jones (LJ) units, with the energy scale *ε*, length scale *σ*, and Boltzmann constant *k*_*B*_ set to unity. The polymer was modeled as a flexible bead–spring chain (Kremer–Grest model) comprising *N* = 800 monomers connected by finitely extensible non-linear elastic (FENE) bonds, with parameters chosen to prevent chain crossing and ensure polymer integrity [46]. Polymer beads and cosolvent particles all had identical diameters *σ*.

Nonbonded interactions between all particles were described by truncating and shifting Lennard–Jones potentials,

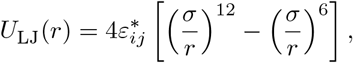

with pair-dependent reduced interaction strengths 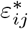. The potential was truncated at a cutoff *r*_*c*_ and shifted to zero at the cutoff. Purely repulsive interactions were implemented by truncating and shifting the potential at its minimum (*r*_*c*_ = 2^1*/*6^*σ*), corresponding to Weeks–Chandler–Andersen (WCA) interactions [47]. Attractive interactions employed a larger cutoff *r*_*c*_ = 2.5*σ*.

Unless otherwise stated, the interaction matrix is specified as follows: polymer–polymer and polymer–P2 interactions are purely repulsive, corresponding to truncated and shifted Lennard–Jones potentials with cutoff *r*_*c*_ = 2^1*/*6^*σ*. Polymer–P1 interactions are attractive with *ε*^∗^ = 1 and cutoff *r*_*c*_ = 2.5*σ*. P1–P1, P2–P2, and P1–P2 interactions are weakly attractive, with 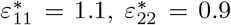, and 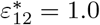, respectively. The chemical potentials are fixed to 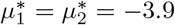. This choice encodes the three key ingredients of PAC, namely attractive polymer–P1 interactions, repulsive polymer–P2 interactions, and weak attraction between P1 and P2. The system was simulated in a cubic box with periodic boundary conditions in all directions.

Equations of motion were integrated using a velocity-Verlet scheme with a time step Δ*t* = 0.0025. All particles were thermostatted using a Langevin thermostat at reduced temperature *T*^∗^ = *k*_*B*_ *T/ε* = 1, providing stochastic thermalization that mimics solvent-mediated friction and noise in coarse-grained biomolecular systems.

To allow the concentrations of the two proteins P1 and P2 to adjust dynamically, both species were coupled to particle reservoirs via grand-canonical Monte Carlo (GCMC) insertion–deletion moves [48] implemented using the LAMMPS fix gcmc algorithm [44]. GCMC updates were invoked every 10^3^ molecular dynamics steps, with 10^3^ insertion–deletion attempts per invocation for each cosolvent species and no additional MC displacement moves. Chemical potentials were specified in reduced form *µ*^∗^ = *µ/ε*, enabling controlled tuning of reservoir concentrations without imposing fixed stoichiometries.

Simulations were initialized with a short soft-potential ramp (10^4^ steps) to remove unfavorable overlaps. The polymer was then equilibrated for 2 × 10^6^ steps in the absence of polymer–P1 attraction and GCMC exchange, while the cosolvent interaction matrix was already present. For each state point, polymer–P1 interactions and GCMC coupling were subsequently switched on, followed by an equilibration run of 6 × 10^6^ steps and a production run of 5 × 10^6^ steps during which configurations were sampled for analysis.

Radial number-density profiles were obtained by computing the polymer center of mass in each configuration and binning particles according to their distance *r* = |**r** − **R**_CM_| into spherical shells of fixed thickness, using minimum-image distances for the cosolvent species. Profiles were averaged over configurations to characterize the spatial organization of polymer, P1, and P2.

## ACKNOWLEDGMENTS

The authors acknowledge support from the Deutsche Forschungsgemeinschaft (DFG) under the grant SO 277/25. H.S. was supported by the Deutsche Forschungsgemeinschaft (DFG, German Research Foundation) under Germany’s Excellence Strategy - EXC-2068 - 390729961. J.U.S. acknowledges support by the DFG under Germany’s Excellence Strategy - EXC2068 - 390729961 - Cluster of Excellence Physics of Life.

